# Young genes to the front - a strategy for future resistance against powdery mildew?

**DOI:** 10.1101/126045

**Authors:** Markus Boenn, François Buscot, Marcel Quint, Ivo Grosse

## Abstract

Nonhost resistance of a plant against a microbial pathogen can be the result of a long-lasting coevolutionary optimization of resource allocation in both host and pathogen. Although this has been suggested for years, coevolutionary aspects leading to nonhost resistance in plants are not fully understood yet. Instead, most studies focus on limited subsets of genes which are differentially expressed in infected plants to describe details of defense strategies and symptoms of diseases.

Here, we exploit publicly available whole genome gene expression data and combine them with evolutionary characteristics of genes to uncover a mechanism of host-pathogen coevolution. Our results suggest that metabolic efficiency in gene regulation is a key aspect leading to nonhost resistance. In addition, we find that progressing host-pathogen coevolution is accompanied by subtle, but systematic overexpression of recently founded genes. In support of our plant-specific data, we observe similar effects in animal species.

## Introduction

Susceptibility of complex organisms to a small number of host-specific microbial pathogens causes diseases and, in plants, significant crop failure and yield losses each year [Oerke et al. 2012]. However, in their natural habitats, organisms are exposed to a much broader range of pathogenic microbes without showing symptoms of disease for the majority of their life cylce. The mechanism responsible for this phenomenon in plants is called nonhost resistance (NHR), the details of its mechanism and evolution are not entirely understood yet [Foley et al. 2013].

NHR is part of the innate immune system of plants. Different from systemic acquired resistance in plants or the adaptive immune system in vertebrates, NHR is not established for only some individuals having been exposed to a certain pathogen. Instead, NHR has been inherited from ancestral plants and affects the entire host plant species [Heath 2000].

A number of studies investigate the plant innate immune system and NHR (reviewed in Ausubel [Ausubel 2005], Jones and Dangl [Jones & Dangl 2006] or Bettgenhaeuser et al. [Bettgenhaeuser et al. 2014], for instance), frequently due to its importance for crop yields. Accordingly, numerous genes and strategies are known to be involved in NHR, causing interaction-specific types of resistance.

Most studies focus on the description of symptoms and the exploration of small subsets of genes, which are significantly modulated in infected plants. However, as NHR is not learned during an individuals life cycle, evolutionary characteristics of genes must provide explanations for presence and absence of NHR. A process describing this phenomon is the arms race [Hulbert et al. 2001]. In the context of hosts and pathogens, the arms race describes host-pathogen coevolution [Jones & Dangl 2006] [Bettgenhaeuser et al. 2014] and is, in addition to coevolution of flowers and their pollinators, probably one of the best studied coevolutionary processes of all living organisms, not only plants.

Immune responses, including NHR, are energetically costly [Segerstrom 2007]. Typically, two types of immune response are distinguished: resistance against a pathogen or disease tolerance. In the latter case, the host focusses on maintainance and repair of the damage caused by the pathogen in order to survive. As suggested by McNamara and Buchanan [McNamara & Buchanan 2005] as well as Segerstrom [Segerstrom 2007], resources consumed for one of the two tasks are not available for other tasks.

Having been established by (co-)evolutionary selection, the interplay of immune response and maintainance should be highly optimized, i.e. ensure survival and ongoing reproduction of the host, while staying efficient in terms of minimal utilization of available resources [Beilharz et al. 1993].

A recently established method to uncover evolutionary trends in entire genomes is phylostratigraphy [Domazet-Lošo et al. 2007]. Using extensive BLAST searches against sequence databases comprising reference protein sequences from a large number of species, this approach is able to assign an approximate evolutionary age to each protein-coding gene of a target organism, based on sequence homology.

In a phylotranscriptomic approach, a combination of phylostratigraphy and gene expression data has been applied to explore various scientific questions, mainly regarding developmental processes like the developmental hourglass in animals [Domazet-Lošo & Tautz 2010], plants [Quint et al. 2012] and also in fungi [Cheng et al. 2015]. Here, amount and complexity of data necessiate to sum up the data to a scalar value. The proposed weighted mean, called Transcriptome Age Index (TAI) [Domazet-Lošo & Tautz 2010], combines gene age and gene expression and is interpreted as the mean evolutionary age of the transcriptome. A complementary measure incorporating information about gene divergence instead of age, the Transcriptome Divergence Index (TDI), has also been proposed [Quint et al. 2012].

In this work we apply phylotranscriptomic methods to investigate how presence of resistance (NHR) differs from absence of resistance. We find that NHR is associated with coevolutionary optimization of the immune response and mainly based on favoured recruitment of older genes. In contrast to this, we further suggest that the host makes significant use of recently founded genes to escape from susceptibility for a host-specific pathogen.

## Material and Methods

### Phylostratigraphy

Phylostratigraphy is a method to estimate the phylogenetic age of the entire set of an organisms genes and is described in detail in [Domazet-Lošo et al. 2007]. For a given target organism, phylostratigraphy splits the tree of life into age classes, the phylostrata. Phylostrata are identified by ps1, ps2, …, psK with psK representing the set of youngest, recently founded, genes and ps1 representing the set of oldest genes, of which domains are conserved in species of all living species. In this work, phylostrata were selected along the lineage of *Arabidopsis thaliana*, according to the NCBI taxonomy database, with ps15 being the set of youngest genes.

Protein-coding genes were assigned to phylostrata using the method of Domazet-Lošo et al. [Domazet-Lošo et al. 2007]. In brief, protein sequences of representative gene models of *A. thaliana* were downloaded from arabidopsis.org (release TAIR10). We used BLAST+ [Camacho et al. 2009] to perform protein-protein searches against the NCBI-NR database (E-value < 1e-5). Retrieved sequences were assigned to a phylostratum, according to the Last Common Ancestor (LCA) of *A. thaliana* and the species the retrieved sequence originates from. Prior to application of BLAST, sequences originating from viruses and environmental samples were removed from NCBI-NR database.

Unless the protein is specific to *A. thaliana*, each query protein has hits in various recent and distant phylostrata, like Arabidopsis, dicots, or eukaryotes. Each query protein was assigned to the most distant phylostratum with a BLAST hit. Query proteins without hit were assigned to the youngest phylostratum ps15.

### Divergence

Estimates of divergence between *A. thaliana* and related species were downloaded from Ensembl-database via biomaRt-package [Durinck et al. 2005] and have been derived by the codeml-function of the PAML package [Yang 1997]. These estimates comprise the number of synonymous substitutions (dS) and nonsynonymous substitutions (dN) per site, of which the ratio dN/dS was calculated. Accorrding to classic evolutionary biology, small dN/dS ratios are indicative of negative selection, whereas large dN/dS ratios are associated with genes under positive selection. However, as the absolute dN/dS value is sufficient for the analyses presented here, it is not necessary to test for specific signatures of selection.

To allow for better comparison between age of genes (represented by discrete phylostrata) and divergence (represented by continuous values), dN/dS ratios were distributed into 5% quantiles (discrete representation), where each 5% quantile is a divergence stratum, identified by ds1, ds2,…,ds20. Here, ds20 represents the set of divergent, fast-evolving, genes and ds1 represents the set of genes being highly conserved in *A. thaliana* and the reference species. In total, 17651 genes were considered for the reference species *Arabidopsis lyrata*.

### Divergence times

Unless otherwise stated, estimates of divergence times between species related to *A. thaliana* and geological times covered by phylostrata were taken from the TimeTree database [Hedges et al. 2006].

### Microarray data & Filtering

We used previously published microarray data (Affymetrix Arabidopsis ATH1 Genome Arrays) investigating NHR in *A. thaliana*. In the experiment the authors challenged plants by two fungal pathogens [Stein et al. 2006]. In the host-specific treatment (H) plants were inoculated with the powdery mildew *Erysiphe cichoracearum* (synonym for *Golovinomyces cichoracearum*). In the nonhost-specific treatment (NH) plants were inoculated with the grass mildew *Blumeria graminis* f.sp. *hordei;* its natural host is barley. Material for four biological replicates per condition (12 samples in total) has been collected from rosettes one day after inoculation. Normalized data was downloaded from NCBI-GEO database (accession GSE3220).

From microarray data only probesets of genes present in the current release of TAIR (TAIR10) were kept. Probesets representing multiple genes were removed. If a gene is represented by multiple probesets (167 genes), expression values of corresponding probesets were summarized by the arithmetic mean.

Together, expression values of 20096 genes were considered for further analyses.

### Additional datasets

To support findings of the main text, we investigated additional datasets based on microarrays and RNA-Seq experiments. For a brief description of design, preprocessing and assignment of phylostrata to genes we refer to Supporting Datasets.

### Regulation strength

We applied fold-change (FC) as a typical measure to assess strength and direction of gene regulation. The FC of each gene was defined as the ratio of mean of raw expression values in treatment and mean of raw expression values in control.

### Regulatome based indices

In analogy to the Transcriptome Age Index (TAI) introduced by Domazet-Lošo and Tautz [Domazet-Lošo & Tautz 2010] (see Supporting Note), the Regulatome Age Index (RAI) was obtained by substituting expression values of each condition in the formula for the TAI by FCs between conditions. The term ‘regulatome’ describes the set of genes modulated in an experiment [Ponomarev et al. 2010]. To enable focussing on the direction of modulation, two types of RAI were defined, one for induction and one for repression.

For a given comparison c and a set of N genes the RAI for up-regulated (induced) genes was defined as

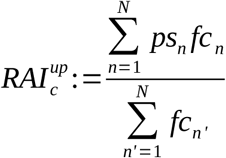
with fc_n_ being the FC of gene n. For down-regulated (repressed) genes, the inverted FC was used:

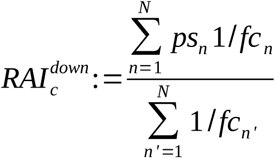

Quint et al. [Quint et al. 2012] introduced a complementary measure, the Transcriptome Divergence Index (TDI), using information about gene divergence instead of gene age. In analogy to the TAI-TDI relationship, phylostrata were substituted by divergence strata in the definition of the RAI. The resulting measure is called Regulatome Divergence Index (RDI). While the RAI provides information about the mean age of gene regulation, the RDI quantifies selective preasure on gene regulation, providing information about selective preasure of modulated genes and possible evolutionary contraints affecting gene regulation.

### Comparison of regulation strengths

To compare the two regulation strengths per phylostratum or divergence stratum, we calculated the difference of 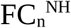 and 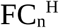, where n is the gene and H and NH are the comparisons. This was done for each direction of modulation separately. The calculated difference can be rewritten as 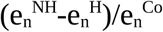, i.e. the difference between expression values, normalized by the expression in control conditions. From resulting stratum-wise distributions, we took the median as a representative value.

Compared to the arithmetic mean, the median has the advantage that it allows for quantifying interpretations, as it divides the set of genes in two groups of equal size. E.g., considering the toy example of gene expression values (1,1,2,2,3,4,4,4,100) we can say that more ‘genes’ have a value greater than the median (m=3), which is not possible for the arithmetic mean (M=13.4) due to the outlier. In the same sense, considering the toy example of differences between gene expression values (-5,-3.5,-4,-2,-1,7,10) we can say that more ‘genes’ have a difference greater than the median (m=-2), which is not possible for the arithmetic mean (M=0.21).

Standard errors for each stratum were obtained by applying a two-sample bootstrap approach within the stratum, given the direction of regulation. In detail, we took a random sample from pairwise differences between treatment-wise FC (with repetition) and calculate the median. This procedure was repeated 1000 times. From the resulting distribution of medians, the standard deviation represents the standard error of the observed median difference between treatments.

### Occupation of metabolic resources

In the absence of an established estimate for metabolic resources used along the entire process of gene expression, from transcription to translation, we used the transcript concentration, i.e. the expression value obtained from microarray data, as an approximation.

Occupation (or release) of resources in treatments with pathogens were estimated by the difference between the expression value in a treatment and the expression value in uninoculated samples.

To assess the amount of expressed transcripts on average, for curves representing host-specific treatment H a segmented regression approach was applied to fit a line to the steady increase accross young genes and to find the point where dominance of resource occupation stops. For the same set of genes a regression line was fitted to the curve representing nonhost-specific treatment.

### Availability of data and methods

To perform analyses, the statistical programming language R was used. Routines were summarized in the R package ‘phyintom’, currently deposited at https://sourceforge.net/projects/phyintom/. The package comprises the routines as well as manuals and a vignette to reproduce essential findings presented in the main text of this work.

## Results

### Choice of data

To understand NHR, it is imperative to also understand what happens when plants are not resistant against a pathogen and are in a stage of coevolutionary optimization. Thus, careful selection of a dataset with a proper experimental design is critical. We choose the dataset from Stein et al. [Stein et al. 2006] with a design as depicted in Figure 1a.

**Fig. 1.**
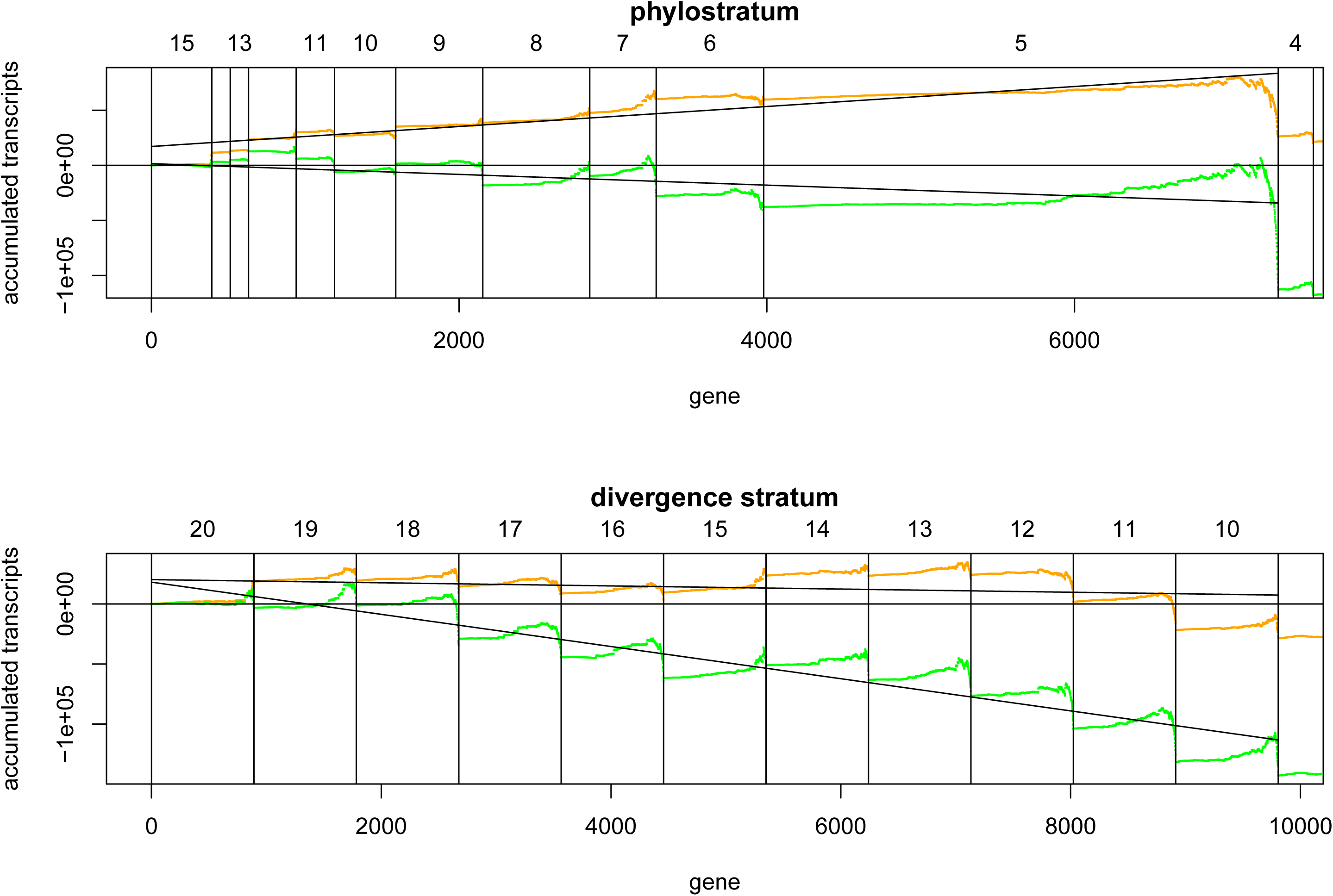
Design of the study. (a): The NHR-experiment considers three groups: an uninoculated control (Co), a group with plants challenged by the host-specific pathogen (H) and a group with plants challenged by a non-host-specific pathogen (NH), which is not compatible to the plant under investigation, but to plants from more distant phlya. (b) and (c): The two treatments are compared to the control group. Genes are considered induced if the fold-change is greater than one (b). Genes are considered repressed if the fold-change is less than one (c). Seven genes exhibit a fold-change of exactly one in the comparison with treatment NH. To ensure that numbers of genes sum up to 20096, they are included in the sets regarding modulation in treatment H.

This dataset covers two essential, mutually exclusive treatments. Plants are inoculated with a host-specific pathogen (H). To compare this state of susceptibility with the opposite state, resistance, plants are treated with a nonhost-specific pathogen (NH) in an independent experiment. Both treatments are compared to control, i.e. uninoculated, samples (Co).

Two directions of modulation are considered. Induced genes are stronger expressed in treatments, compared to the control (Figure 1b). Repressed genes are stronger expressed in control, compared to the treatment (Figure 1c). The large overlap between comparisons indicates high agreement of both treatments, considering only the direction but not the strength of modulation.

### Age and divergence correlate with expression and regulation

To determine phylostrata [Domazet-Lošo et al. 2007], protein-coding genes of *A. thaliana* were assigned to a set of 15 distinct phylostrata (see Table 1), each representing the evolutionary age of a certain set of genes. According to previous findings [Wolf et al. 2009], we confirm that expression values of these genes increase with age (Supplementary Figure S1).

**Table 1:** Phylostrata along the lineage of *A. thaliana*

| ps | taxon | genes |
| --- | --- | --- |
| 15 | Arabidopsis thaliana | 393 |
| 14 | Arabidopsis | 119 |
| 13 | Camelineae | 119 |
| 12 | Brassicales | 310 |
| 11 | rosids | 249 |
| 10 | dicots | 398 |
| 9 | Mesangiospermae | 566 |
| 8 | Magnoliaphyta | 695 |
| 7 | Spermatophyta | 432 |
| 6 | Tracheophyta | 699 |
| 5 | Embryophyta | 3344 |
| 4 | Streptophyta | 228 |
| 3 | Chlorobionta | 647 |
| 2 | eukaryotes | 5996 |
| 1 | cellular organisms | 5901 |
|  | total | 20096 |

We used the log_2_-transformed fold change (log(FC)) to create a plot in analogy to Supplementary Figure S1 for gene regulation, shown in Supplementary Figure S2. As the FC provides information about the direction of the modulation, phylostratum-wise distributions of transformed FC are shown for induced and repressed genes separately.

We find that induction of genes systematically decreases with gene age, i.e. the younger the genes are, the larger is their FC. This is observed for both treatments, but the relationship is stronger for treatment H. For repressed genes, no systematic dependence on age is visible for treatment H (see Supplementary Table S1), but for treatment NH. Here, Kendall’s rank-based correlation coefficient indicates stronger repression of young genes.

It is possible, however, that correlations between age and magnitude of modulation are artificial. Gene expression data used in this work has been normalized using MAS 5.0 for Affymetrix microarrays [Stein et al. 2006]. The FC calculated from such data tends to be biased towards larger values when low transcript concentrations are involved, compared to a FC from high transcript concentrations [Wu et al. 2004]. Hence, recalling that young genes exhibit low expression values (Supplementary Figure S1), slopes shown for induced genes are less surprising.

From a technical point of view, these observations have significant impact on further analyses. Typically, 1.5- or 2-fold modulation of genes is considered to be meaningful. Applying these cutoffs to the data (horizontal gray lines in Supplementary Figure S2), a large proportion of genes assigned to distant (old) phylostrata would be excluded from further analyses.

To analyse the relationship between expression values and selection pressure on genes as well, we assigned dN/dS ratios (in the context of this study synonymously used with the term *sequence conservation*) to 20 divergence strata. According to previous studies [Quint et al. 2012] [Drost et al. 2015] we confirm that age and sequence conservation exhibit only weak dependence according to Kendall’s rank-based correlation coefficient and Cramer’s V (Supplementary Figure S3), suggesting that they are complementary measures for evolutionary studies.

We find that gene expression is not independent from sequence conservation (Supplementary Figure S4). In all conditions, very conserved genes exhibit high expression values. This is in line with the suggestion of Drummond et al. [Drummond et al. 2005] that highly expressed genes evolve slowly to avoid cytotoxic protein misfolding.

In analogy to Supplementary Figure S2, we also explore the relationship between FC and sequence conservation. For treatment with H we find that conserved induced genes (low divergence strata) exhibit lower FCs, while no dependence on sequence conservation for repressed genes can be detected (Supplementary Figure S5, Supplementary Table S1). Dependence on sequence conservation in treatment NH is significantly weaker for induced genes and inidicates that conserved genes exhibit lower FCs. In contrast to this, repression affects conserved genes only mildly in this comparison.

Despite the possibility of a bias introduced by microarray normalization, regarding dependence on age and divergence there are distinct differences between treatments. These differences are likely to be of biological rather than technical origin. However, based on these observations we can not apply one of the traditional cutoffs (or any other global cutoff). Instead, we consider induced genes as genes having FC>1 and repressed genes as genes having FC<1.

### Resistance is achieved efficiently

Induced defenses are accepted as effective strategies of plants to fight against attacks by herbivores and pathogens [Karban & Myers 1989] [Kessler & Baldwin 2004]. Even more, in terms of bioenergetics induced strategies for resistance are suggested to be cost-saving as they are not activated when resistance expression is not required [Karban et al. 1997].

Hence, following the cost-benefit paradigm, resistance against the nonhost-specific pathogen NH should be efficient and achieved at minimum usage of resources. In this case the number of induced genes should be much smaller when the plant is treated with NH than with H. Otherwise, the plant takes similarly high efforts in presence of H although immune response is insufficient and ineffective.

Figure 1b reveals induction of numerous genes in both treatments, contradicting a constitutive defense strategy. We find that the number of genes induced in the presence of the nonhost-specific pathogen NH (N=8710, Figure 1b) is significantly smaller than one would expect by chance (Binomial test, P<2e-16). In contrast to this, the null hypothesis of similar numbers of induced and repressed genes cannot be rejected when the plant is inoculated with pathogen H (N=10145, P=**0.17**). Accordingly, the number of genes induced in NH is significantly smaller than the number of induced genes in H (McNemar’s test, P<2e-16).

Vice versa, Figure 1c trivially reveals repression of large numbers of genes in presence of NH. This supports the idea of a cost-saving immune response and is in line with findings of previous NHR studies, focussing on much smaller sets of genes [Zimmerli et al. 2004] [Stein et al. 2006] [Foley et al. 2013].

### Regulatome based indices reveal a general pattern of NHR

To get a general evolutionary pattern describing NHR in plants, we applied two regulation based indices, the Regulatome Age Index (RAI) and the Regulatome Divergence Index (RDI). These indices combine relative gene expression and gene age/conservation and focus on gene induction and repression. They outperform typical transcriptome based indices like TAI and TDI under several aspects (see Supporting Note).

Both directions of modulation were taken into account. Accordingly, Figure 2 shows results for RAI^up^ and RAI^down^ as well as RDI^up^ and RDI^down^. Considering gene induction, we find that modulated genes are less susceptible to evolutionary changes in treatment NH (red lines). Here, induction affects most notably older genes and genes under weak selection (low values of RAI and RDI, respectively), compared to treatments with the host-specific fungus H. Vice versa, repression affects older genes and genes with high dN/dS ratios when the plant is inoculated with H. Reliability of observed differences between comparisons is indicated by standard errors and confirmed for the RAI by a z-test (P<0.01 for induction as well as for repression). For the RDI, differences between comparisons are not significant (RDI^up^, P<0.11) or only weakly significant (RDI^down^, P<0.05), respectively.

**Fig. 2.**
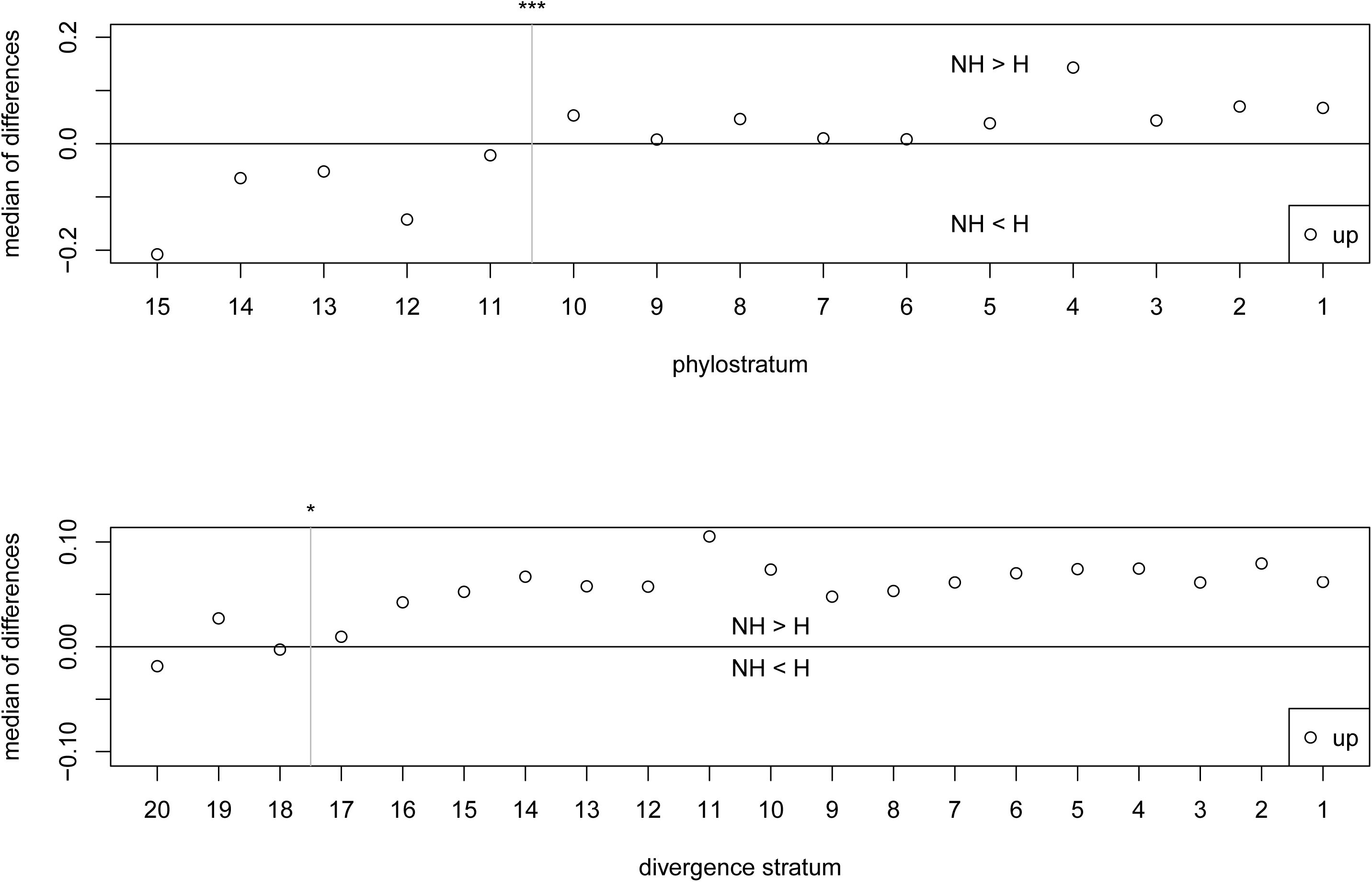
Regulatome Age Index (RAI) and Regulatome Divergence Index (RDI) for the data. Subfigures (a) and (b) represent RAI and RDI, respectively. The RDI is given for the related species *Arabidopsis lyrata*. Measures distinguish between the two directions of modulation, RAI^up^ (or RDI^up^) and RAI^down^ (or RDI^down^) for induction and repression of genes, respectively.

For a fair comparison with the TAI, the RAI was calculated accross all genes. However, considering subsets of genes is more intuitive. Accordingly, Supplementary Figure S6 shows RAI profiles for (1) taking into account only all induced genes (10139 for H, 8714 for NH), (2) 6906 commonly induced genes, as well as (3) genes which are exclusively induced in each treatment (3229 for H, 1808 for NH) and harbour genes which are very specific to the corresponding type of interaction. For repressed genes RAI values are calculated for analogous subsets.

Although the dominating shape of the ‘full’ RAI profile is not changed in any case, consideration of subsets of genes clearly increases significance of observed differences in cases (1) and (3). However, reliability of indicated differences between comparisons in case (2) is weak, in particular for commonly repressed genes. Together, Supplementary Figure S6 indicates that the NHR pattern is mainly caused by the relatively small number of genes which are highly specific to the type of interaction, being modulated to the opposite direction in the other treatment (see Figure 1).

### Regulation strengths of commonly modulated genes

Although observed differences between treatments are not significant in RAI profiles, due to their sizes sets of commonly modulated genes are likely to harbour not only noise, but genes being relevant for both types of treatments. As immune responses can be expected to be induced, we focus on commonly induced genes first.

We want to investigate, if specific phylostrata ranges are responsible for the shape of the RAI profile. For this, we calculated gene-wise differences of regulation strengths as expressed by FCs. From the resulting distribution we take the median as a representative value. This is done for each phylostratum. The median allows to draw conclusions about the number of genes exhibiting higher or lower induction in one of the treatments, hence combining number of induced genes and strength of induction. Medians are greater than zero, when modulation is stronger and affects more genes in the nonhost-specific treatment. Otherwise, medians are less than zero.

NHR is complex due to the involvement of numerous pathways [Gill et al. 2015], requiring strict control of gene expression by likewise complex regulatory mechanisms, which are most notably observed for old genes [Warnefors & Eyre-Walker 2011]. Accordingly, Figure 3a reveals that strength of induction is systematically higher for old genes when considering plants treated with NH. By construction of phylostrata [Domazet-Lošo et al. 2007], functions of expression products of these genes tend to base on evolutionarily optimized domains without significant modifications since their first appearance. In contrast to this, plants treated with the host-specific pathogen H systematically exhibit stronger induction of younger, recently founded genes. Together, this results in a sigmoidal arrangement of phylostratum-wise medians.

**Fig. 3.**
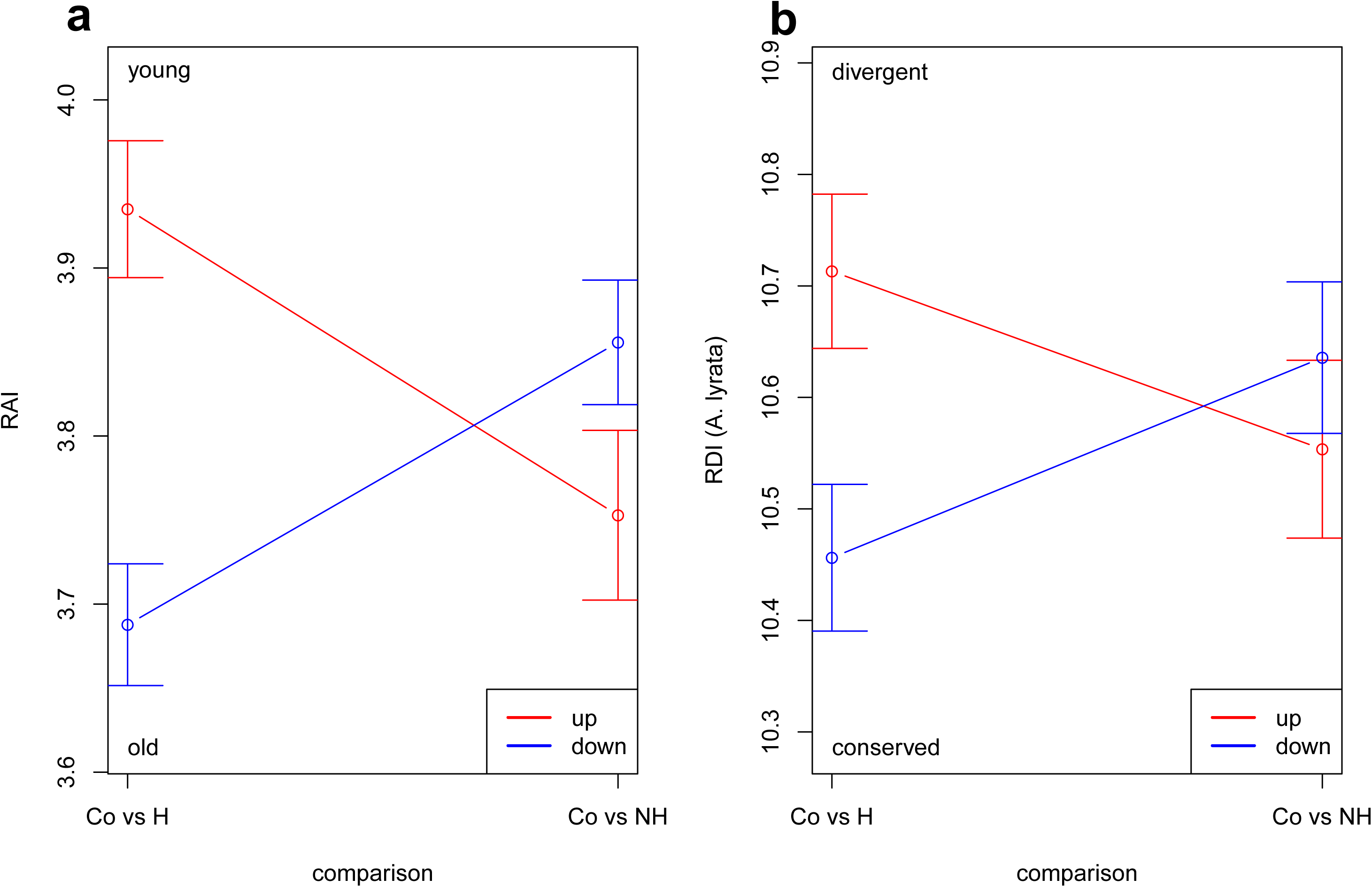
Comparison of commonly induced genes. Medians of pairwise differences of genes in each phylostratum and divergence stratum are shown for commonly induced genes. Horizontal zero lines determine whether regulation in treatment NH (points are above line), or treatment H (points are below line) is stronger. Strata comprising young or fast-evolving genes, respectively, are located on the left. Vertical gray lines seperate points at right-most stratum having points below zero line, allowing for a single outlier. Stars indicate significance of enrichment of points below zero line (Fisher’s test, *: P<0.05, ***: P<0.001).

Applying the same procedure to divergence strata as well (Figure 3b) reveals weak evidence that genes under strongest selective pressure are stronger induced in treatment H, while conserved genes tend to be stronger induced in NH.

Supplementary Figure S7 reveals that most commonly repressed genes are stronger repressed in presence of NH, no matter the stratum.

### Systematic induction of young genes

Results obtained so far were derived from FCs (Figure 2) and differences between FCs (Figure 3). They suggest treatment-specific favour of age ranges regarding strength and direction of modulation by condensing data or consideration of a large subset of genes.

However, the FC provides information about relative changes in transcript concentrations. Hence, it is unable to distinguish between inductions from, e.g., 1 to 10 and 100 to 1000 transcripts (10-fold in both cases). To further understand how absolute changes in transcript concentrations affect NHR, we next consider absolute differences between transcript concentrations of control and treatment groups.

Using absolute differences has the advantage that also information in terms of occupation of resources is provided; in this case numbers of transcripts serve as an approximation for resources.

For a comprehensive view on the data, genes were sorted according to their phylostratum, young genes first. Within each phylostratum, genes were sorted by transcript concentration in control conditions, lowest values first. Then, differences between treatment and control were cumulatively added to each other, accross all genes, beginning with young genes. This is motivated by the finding that large numbers of young genes exhibit distinct and treatment-specific behaviour, as they are strongest induced in H (Figure 3). When values increase, induction is indicated. Otherwise, genes are repressed. To focus on young genes, curves covering recent phylostrata are shown in Figure 4. Curves for the entire dataset are given in Supplementary Figure S8.

**Fig. 4.**
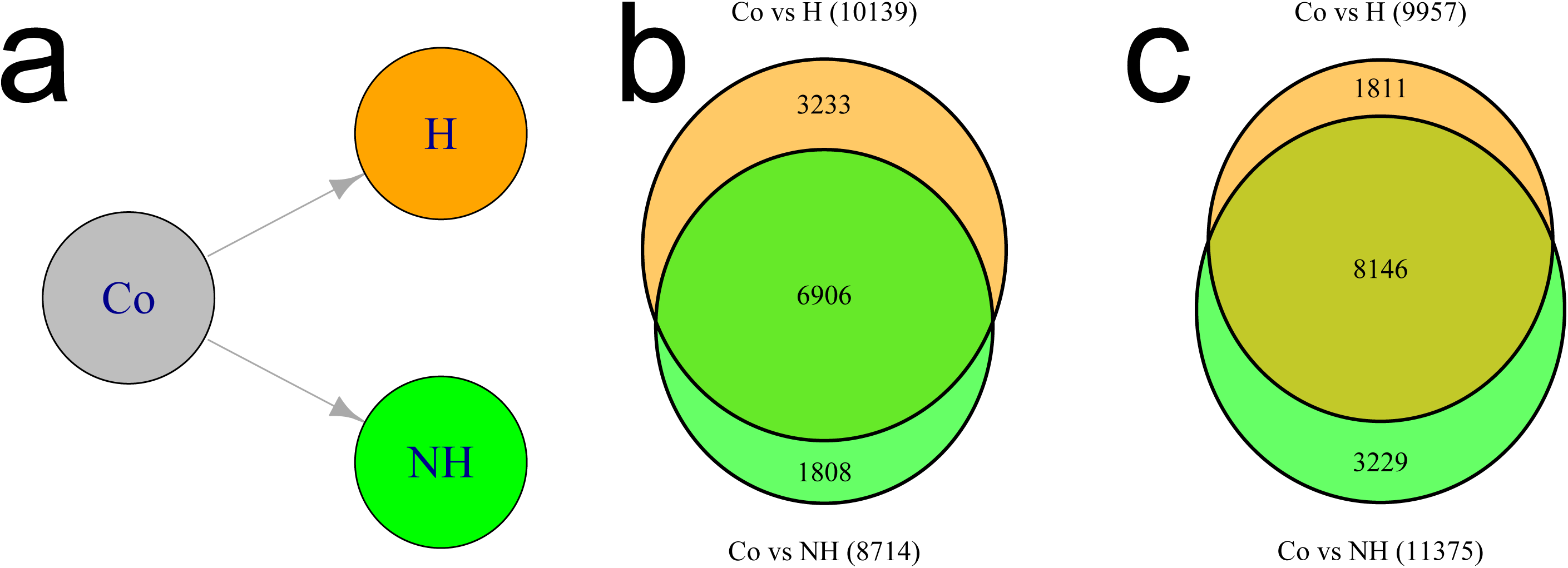
Accumulation of transcript expression accross genes. Figure shows cumulative curves of differences between expression levels in control and treatment groups. For this, genes are ordered by their phylostratum (or divergence stratum), so that young (divergent) genes are on the left and old (conserved) genes are on the right. Borders between strata are given by vertical lines. Within each stratum, genes were sorted according to their expression value in uninoculated plants (lowest first). Each point corresponds to a gene. Curves are given for comparison with host-specific treatment H (orange points) and nonhost-specific treatment NH (green points). When genes are induced or repressed, the curve increases or decreases, respectively. Curves are given for youngest (most divergent) genes only.

While results obtained so far suggest that numerous young genes are stronger induced in H, the immediate and steady increase of the orange curve in Figure 4 suggests that in fact all young genes either (i) are induced in H or (ii) do not experience significant repression. Extending the conclusion that young genes are stronger induced in H (Figure 3), this systematic induction not only comprises younger phylostrata ps15 to ps11, covering 1190 genes, but reaches back even to the evolutionarily old phylostratum ps5 (Embryophyta), covering 7324 genes in total (36.4% of all genes). Indeed, 3872 young genes in these phylostrata exhibit a FC>1.

To get an impression of the number of transcripts added by each gene, a regression line was fitted to the curve for genes contained in ps15 to ps5. We find that, on average and compared to the control, gene expression increases by nine (rounded from slope=9.08, R^2^=0.86) transcripts per gene in H.

In contrast to this, the slope of the curve representing NH for the same range of phylostrata points to the opposite direction, revealing that gene expression decreases by five (-4.87, R^2^=0.41) transcripts per gene. Indeed, only 2854 young genes exhibit a FC>1.

From these observations we extract two pieces of information. First, young genes are less likely to play important roles in an induced immune response against NH. Second, in a cell with limited resources (in terms of free nucleotides, for instance), compared to control conditions young genes release resources, which are likely reallocated to increase expression of old genes.

The curves furthermore suggest that very highly expressed genes (located at right borders of phylostrata) are repressed in each treatment. This is, however, an artifact caused by the arrangement of data points. Genes exhibiting a very high expression value in control condition are outliers in the corresponding stratum and are likely to exhibit a much smaller expression value in any treatment. This consistently results in systematic drops at the end of each stratum.

To confirm that this does not affect our findings significantly, we recreate Figure 4 without sorting genes according to their transcript concentration. Instead, within each phylostratum genes are randomly permuted, followed by computation of the cumulative curve. This procedure is repeated ten times and the mean cumulative curve is considered (Supplementary Figure S9). We find that young genes still exhibit an immediate and steady increase in H, which is not visible for NH.

However, the slope stops at the slightly younger phylostratum ps6 (Tracheophyta). This also affects the number of transcripts expressed on average, now being about 14 for H, with R^2^>0.95. Permutation also affects the curve representing NH. Here, the negative trend is increased, i.e. more negative, resulting in about ten transcripts for which no resources in terms of free nucleotides or ribosomes have to be occupied (R^2^>0.85).

We applied the same analyses for divergence strata as well (Figure 4 and Supplementary Figure S8). We find that treatment with NH results in immediate and steady decrease of the curve, indicating that significant repression of genes dominates this treatment. In contrast to this, the 50 % of genes being under lowest negative selection (meaning dN/dS close to one) are dominated by induction, when the host-specific treatment is considered. This is indicated by the curve for H, which is located above the zero line for divergence strata ds20-ds11. For the remaining 50 % of genes being more conserved, repression dominates. Again, this general impression does not change by permutation and averaging (Supplementary Figure S10).

### Additonal datasets

Application of the same approach to a second NHR experiment (again A. thaliana challenged with H and NH fungi, see Supporting Datasets for details) confirmed most patterns derived for the experiment of Stein et al. [Stein et al. 2006] (see Supplementary Figure 11 and 12). However, dataset-specific differences are visible. E.g. the NHR pattern derived by the RAI is not significant. For commonly induced genes, consideration of divergence strata reveals that more weakly conserved genes are involved in interaction with H. Furthermore, systematic resource occupation does not affect fast-evolving genes in H, but in NH.

We also investigated a dataset dealing with rice as well as datasets dealing with animal hosts, which are designed in a fashion that is comparable to Figure 1a (see Supporting Datasets for details). For this, again we assigned genes to phylostrata using the method introduced by Domazet-Lošo et al. [Domazet-Lošo et al. 2007]. Numbers of genes per phylostratum can be found in Supplementary Table S2. Next, we computed the RAI and accumulated transcript concentrations for each species. Considering Supplementary Figure 13 to 16, we find that in treatment H young genes accumulate transcripts in these datasets as well, mirroring the findings of Figure 4. However, the general NHR pattern exhibited by the RAI for A. thaliana (Figure 2) is visible in only some cases. Interestingly, in particular the NHR pattern is not visible when considering mice (Supplementary Figure 15 and 16), which rely on both innate and an evolutionarily much younger acquired immune system [Zhu et al. 2013].

## Discussion

Applying phylotranscriptomic methods, we have observed a systematic activation of thousands of young genes during a compatible interaction between a host and a microbial pathogen H. Moreover, we observed that this activation is specific to the compatible interaction. In contrast to this, during an incompatible interaction (NH) recruitment of old genes is favoured.

We presume that activation of thousands of young genes is a sophisticated coevolutionary strategy of the host and a key element of the arms race. Here, functions of induced young genes, which, by construction of phylostrata [Domazet-Lošo et al. 2007], harbour previously not established domains, are combined with functions of induced old genes. This might generate new ways for detection of microbial effectors and proper responses, lowering susceptibility for the pathogen.

However, as complex regulatory mechanisms are rare for young genes [Warnefors & Eyre-Walker 2011], a directed activation of large amounts of young genes appears to be unlikely. Instead, from our point of view the results propose an undirected and trial-and-error-based strategy (TES) of the host.

Young genes are usually short and consist of a low number of exons. This has been found for animals by Neme and Tautz [Neme & Tautz 2013] and can be confirmed for A. thaliana (Supplementary Figure S17 and S18). Further, their regulation requires fewer transcription factors and they harbour lower numbers of other regulatory and structural elements [Warnefors & Eyre-Walker 2011]. These characteristics of young genes indicate rapid transcription and posttranscriptional processing. Subsequently, initiation and elongation during translation of short genes tend to be faster [Ding et al. 2012]. Hence, irrespective of their originally intended biological function, expression products of young genes are rapidly available for the immune response.

In addition to this and in line with the argumentation of Drummond et al. [Drummond et al. 2005], undirected induction of young genes is less risky than undirected induction of old genes, which is a further benefit. As recently founded genes tend not to be involved in complex regulatory pathways [Warnefors & Eyre-Walker 2011], their induction is rarely expected to accidently have negative impacts on well established and essential pathways controlling growth and metabolism, for instance.

On the other hand, the systematic induction of thousands of young genes is a metabolic challenge. The biosynthesis of nine transcripts and proteins on average occupies significant amounts of resources, ranging from free nucleotides and RNA polymerases for transcription to free ribosomes and t-RNAs for translation. As resources in cells are limited and genes are competing for them [Brewster et al. 2014], they have to be reallocated towards younger genes. Vice versa, when used for this task, they cannot be used for other tasks [Segerstrom 2007] [McNamara & Buchanan 2005], e.g. expression of old genes. Hence, the observed slightly lower induction of old genes (Figure 3) might, at least in part, be a consequence of the systematic induction of young genes.

An obvious contradiction in this scenario is that the host induces young genes at the cost of old genes, potentially lowering the effectiveness of the part of the immune response against H, which is based on old genes. However, tolerating disease by an only partially efficient immune response can be sufficient to survive pathogen attack and has been suggested to increase fitness of the host [Rauw 2012]. At the coevolutionary stage of susceptibility the host is lacking mechanisms to detect and respond to all effectors elicted into the cell by the pathogen. Hence, with an alternating arms race in mind, we suggest that the TES is an investment into future defense strategies and is to be prefered over investment of too many resources for an unpromising defense response.

We analyzed datasets from additional independent studies involving compatible interactions (like H) and incompatible interactions (like NH) to confirm the presence of a TES. Here, we took into account a second dataset dealing with the host A. thaliana, and datasets dealing with *Oryza sativa* and the animal hosts *Mus musculus* and *Caenorhabditis elegans*.

Beside the diversity of hosts representing two eukaryotic kingdoms, setups of these experiments are highly heterogenic regarding the utilized high-throughput plattform (microarrays and RNA-Seq) and preprocessing of data as well as types of pathogens (bacterial, eukaryotic and viral) used.

Surprisingly, although these and other differences usually result in lower comparability between experiments and agreement in their outcomes [Ingersoll et al. 2010] [Wang et al. 2014] [Maboreke et al. 2016], we repeatedly find that curves representing H exhibit a steeper positive slope accross significant amounts of young genes, compared to curves representing NH.

This similar overall logic of the immune response supplements other examples of convergent evolution towards similar components of the immune systems of plants and animals [Ausubel 2005], indicating a re-invention of the same idea to overcome susceptiblity for a challenging pathogen.

Important questions remain as to how the host initiates and controls the undirected expression of young genes during a TES. Phylotranscriptomic methods provide the opportunity to apply powerful analyses to address these questions, provided the availablity or generation of proper datasets.

## Conclusions

In this phylotranscriptomic study we explored publicly available data and propose that successful resistance of a plant against a nonhost-specific pathogen is caused by efficient use of old genes, indicating that NHR is less susceptible to evolutionary changes. We uncovered a potential approach how coevolution between pathogens and hosts works and found hints that this mechansim is also established in animals, indicating a re-invention of the same idea accross eukaryotic kingdoms.

In this work we analysed data of the host. However, it is possible, that microbial pathogens similarly rely on induction of recently founded genes to overcome defense strategies of the host. This will be subject of further investigations.

Taken together, our results supplement currently existing knowledge about evolutionary aspects of NHR and coevolution of hosts and pathogens.

## Acknowledgments

We thank Robert Paxton for helpful comments on animal hosts. This work was partially supported by a Flexible Pool Grant (number 50170649_#20) of the German Centre for Integrative Biodiversity Research (iDiv) Halle – Jena – Leipzig.

